# KNOB K180 CONSTITUTIVE HETEROCHROMATIN OF MAIZE EXHIBIT TISSUE-SPECIFIC CHROMATIN SENSTITIVE PROFILES DISTINCT FROM OTHER TYPES OF HETEROCHROMATINS

**DOI:** 10.64898/2026.04.01.715864

**Authors:** Mariana C Sattler, Ayush Singh, Hank W Bass, Mateus Mondin

## Abstract

**Background:** Maize knobs are regions of constitutive heterochromatin that are readily identified in both meiotic and somatic chromosomes. These structures have been characterized as stable throughout the cell cycle, exhibiting late replication during the S-phase, and are composed of two specific families of highly repetitive DNA sequences: K180 and TR-1. Although widely used as cytogenetic markers due to their variability in number and chromosomal position across inbred lines, hybrids, and landraces, little is known about their chromatin structure and dynamics. In this study, we analyzed chromatin accessibility of knobs using DNS-seq data across four maize tissues representing distinct developmental stages.

**Results:** Our results reveal that K180 knobs exhibit tissue-specific variation in chromatin accessibility, transitioning between open and closed states during development. In contrast, the TR-1 knob of chromosome 4 remained consistently inaccessible across all tissues analyzed. A knob composed of both K180, and TR-1 further supported this observation, with only the K180 region showing dynamic accessibility. To validate these findings, we also analyzed other repetitive regions such as centromeres, which showed a uniformly closed chromatin structure similar to TR-1. These results suggest a unique developmental modulation of chromatin accessibility associated with K180 repeats. While the chromatin accessibility of knobs does not reach the levels observed at Transcription Start Sites (TSS), the comparison among different classes of repetitive DNA within maize constitutive heterochromatin provides compelling evidence for sequence-specific and tissue-specific chromatin dynamics.

**Conclusions:** Our findings uncover a previously unrecognized property of maize knobs and establish a reference for future studies on chromatin organization and epigenetic regulation of repetitive DNA in plant genomes.

## Introduction

Constitutive heterochromatin was first characterized by Heitz (1928) as densely stained regions of compacted chromatin within the cell nuclei of mosses throughout the cell cycle. Unlike the contrasting less compacted euchromatin, heterochromatin is generally less genetically active and often composed of satellite DNA or highly repetitive sequences. This was confirmed by the isolation and hybridization of satellite DNA to heterochromatin regions (Pardue and Gall 1970). Additionally, the description of the repetitive DNA sequences as a major but largely inert component of the genome (Britten and Kohne 1968) aligned with the initial speculation about constitutive heterochromatin made by Heitz. However, decades of research since then have slowly overturned this notion of heterochromatin as genetically inactive or biologically irrelevant junk DNA.

From studies in animals and plants, we find many cases where the constitutive heterochromatin may contain within them active domains with essential genes expressed during development (Dimitri et al. 2005). For instance, in humans and mice, satellite DNA repeats in heterochromatin can be overexpressed in epithelial cancers (Ting et al. 2011). In *Arabidopsis*, pericentromeric 180-bp repeats and *Athila* retrotransposon respond to pathogen attack by a reduction the silencing mark, cytosine methylation, and a loosening of chromocenter heterochromatin in interphase (Pavet et al. 2006). Transcriptome analysis has shown that constitutive heterochromatic regions contribute significantly to the cellular non-coding RNA pool, an essential part of the multi-layered regulatory network (Lakhotia 2017). Also, variation in the amount of constitutive heterochromatin per genome changes between and within species, including genomic copy number variation coupled to environmental factors such as habitat, ecology, and altitude (Vosa 1996). These exceptions to the general rule of genetically inert heterochromatin highlight the fact that amongst the background of repression and silencing, there are many contexts in which heterochromatin can exhibit dynamic and active properties.

In the maize chromatin, constitutive heterochromatin is conspicuous in two different types of chromosomal regions, the centrally positioned pericentromeric regions and the more distal or terminal structures called knobs. Since their usage by B. McClintock (McClintock 1930) to study transmission genetics in maize, knobs remain as one of the most captivating features of the maize genome. The heterochromatic knobs are so-called because of their small discrete spherical structures along the otherwise fiber-like meiotic chromosomes. They undergo late replication during the cell cycle, are primarily comprised of tandem repeats, and provide clearly visible structures for cytogenetic analyses with light or fluorescence microscopes (Pryor et al. 1980; Aguiar-Perecin et al. 2000; Dawe and Hiatt 2004; Mondin et al. 2014; Bass et al. 2015). Unlike other tandem repeats that occur in strikingly low recombination regions, such as centromeres and telomeres, knobs manifest as multi-megabase blocks of tandem repeats situated in mid-arm and terminal regions of maize chromosomes and they can possess neo-centromeric activity and non-Mendelian meiotic drive in certain genetic backgrounds (Dawe and Hiatt 2004). The potential locations of knobs in the chromosomes are relatively fixed in maize, but within any isogenic inbred line, there exists heritable variation in which specific subset (number) of knob locations occur and the sizes (small or large) at those locations. The same is true for different maize races and varieties, establishing them as valuable genetic markers (McClintock et al. 1981; Aguiar-Perecin et al. 2000; Ananiev et al. 2000; Adawy et al. 2004; Ghaffari et al. 2013; Mondin et al. 2014; Carvalho et al. 2022).

At the DNA sequence level, knobs consist of arrays of tandemly repeated sequences belonging to one of two major classes, K180, made up of the classical and most common 180 bp knob repeat, and TR-1, made up of a 350 bp repeating sequence (Peacock et al. 1981; Ananiev et al. 1998a). A given knob might be exclusively composed of one family or by a mixture of both (Peacock et al. 1981; Ananiev et al. 1998a; Kato et al. 2004; Ghaffari et al. 2013). In cases where both types of repeats coexist within the same knob, they appear to exist in knob-type clusters within the larger knob region (Hiatt et al. 2002). The proportion of K180 repeats on the genome and the number of knobs composed of K180 repeats are usually higher than TR-1 in most maize lines (Hiatt et al. 2002; Kato et al. 2004; Yang et al. 2013). Interestingly, quantitative filter-blot hybridization, FISH, and whole genome sequencing has shown that not all occurrences of these tandem repeats, especially the lower copy number clusters, appear as cytologically distinct structures. Consequently, there exists a semantic distinction between cytological versus molecular knobs, which reflects the cellular-level chromosomal knobs versus the molecular level knob repeat DNA sequences, respectively.

When it comes to their origin and roles, maize knobs pose as puzzling structures. They have been correlated with different genetic effects, including influence on recombination frequency (Ghaffari et al. 2013; Stack et al. 2017), neocentromeric activity and meiotic drive (Rhoades and Vilkomerson 1942; Hiatt et al. 2002; Dawe 2022), correlation with the phylogeographic distribution of different maize populations (McClintock et al. 1981), as well as with relevant phenotypic traits (Pierozzi et al. 1997; Carvalho et al. 2022). The flowering time, for instance, has been correlated with the presence/absence of specific knobs (Chughtai and Steffensen 1987; Carvalho et al. 2022).

In terms of chromatin organization, nucleosome occupancy and density is generally similar between heterochromatin and euchromatin in flies and humans (Chereji et al. 2019). Nucleosome occupancy and distribution is one of the features of chromatin structure that is related to the chromatin condensation levels, gene expression, genome organization within nucleus, faithful segregation during cell division, as well as transcriptional silencing of repetitive DNA sequences (Talbert et al. 2019). To study nucleosome dynamics across the genome, Differential Nuclease Sensitivity sequencing (DNS-seq) has been shown to be useful in maize (Vera et al. 2014; Turpin et al. 2018). This method involves digesting DNA with Micrococcal Nuclease (MNase) under both light and heavy conditions, resulting in DNA fragments that are used to construct library pairs for sequencing. The aligned library reads provide insights into the sensitivity or resistance of specific genomic regions to nuclease digestion. Regions with differential sensitivity delineate open versus closed chromatin, where open chromatin is characteristic of active genic regions, and closed chromatin is more resistant and characteristic of inactive or heterochromatic regions. Consequently, DNS-seq is an effective tool for assessing the structural and functional dynamics of chromatin (Vera et al. 2014; Rodgers-Melnick et al. 2016; Turpin et al. 2018).

DNS-seq profiling of four reference tissues from B73 maize (*Zea mays* L.) is publicly available (Turpin et al. 2018). These chromatin profile datasets are also potentially ideal for examining non-genic and repetitive DNA regions, such as knobs. However, no studies have used this resource to investigate heterochromatin for tissue-specific analysis. We leveraged these chromatin profiling data to investigate the previously unexplored chromatin accessibility and sensitivity for the K180 and TR-1 knob regions in maize. Here, we present our findings, which reveal that among various classes of repeats, the K180 sequences within heterochromatin knobs display distinct and previously unreported tissue-specific chromatin structural features.

## MATERIAL AND METHODS

All data used in this work were obtained from Genomaize (http://www.genomaize.org), a FSU UCSC genome browser for *Zea mays* developed by Daniel Vera, Katie Kyle and Hank W. Bass.

### Chromatin accessibility in knobs

The first step of this research was the visual inspection of chromatin structure accessibility in knobs composed of K180 and TR-1 repeats in the genome of *Z. mays* B73v5. Chromatin accessibility was accessed by DNS-seq data (Vera et al. 2014; Turpin et al. 2018) available on Genomaize as a browser-ready data track. DNA-seq is based on the digestion of the DNA by micrococcal nuclease (MNase) in two digest conditions, “light” and “heavy”. The resulting DNA fragments are quantified by Next Generation Sequencing to obtain the relative abundance of aligned fragments. The difference between the “light” and “heavy” relative read coverage is presented as Differential Nuclease Sensitivity (DNS) values. Therefore, the DNS values of a genomic region might be positive, indicating more accessible chromatin (sensitive to MNase digestion), or negative in regions with more closed chromatin (resistant to MNase digestion). DNS-seq profiles were available for four different tissues: endosperm 15 days after pollination (EN), 1 mm terminal root tips (RT), 1–2 cm earshoots (ES), and 4–5 day old seedling coleoptile nodes (CN) (Vera et al. 2014; Turpin et al. 2018).

The annotated positions of the K180 and TR-1 repeats in the *Z. mays* B73v5 genome were acquired from the TE B73v5 NAM Repeats track (Hufford et al. 2021). This track encompasses annotations for the repetitive fraction of the *Z. mays* genome, including mobile elements and tandem repeats. The track file was downloaded from Genomaize and uploaded to the public Galaxy server (https://usegalaxy.org/, The Galaxy Community 2022). Using the Filter tool, all the K180 (TE;knob180;EDTA;knob) and TR-1 (TE;TR-1;EDTA;knob) knob repeats were filtered and saved to individual BED files. In addition to knob repeats, the positions of CentC repeats were also filtered and saved in a separate BED file. The BED files for each tandem repeat family (K180, TR-1 and CentC) were then reloaded into the Genomaize.org genome browser as individual tracks, allowing us to inspect their DNS-seq patterns separately.

### Z-score-based comparison of knob chromatin accessibility in four tissues

The statistical comparison of knob DNS-seq patterns among the different tissues was based on z-scores. For this, DNS-Seq data of the four tissues (CN, EN, ES and RT) were downloaded from Genomaize and used for calling positive peaks (MNase hypersensitive sites) with the MACS3 algorithm (Zhang et al. 2008) using the following parameters: q-value of 0.05, nomodel, fragment length as minimum peak size and maximum gap.

The positive called peaks, corresponding to accessible chromatin sites within each tissue, were used to generate a probability distribution of expected intersections for each repeat family using BEDTools (Quinlan and Hall 2010). Initially, the positions of positive called peaks were shuffled, and those new positions were intersected with each of the repeat families. This process was repeated one hundred times for each repeat family (K180, TR-1 and CentC) within each tissue.

Based on the probability distribution, a z-score was calculated to determine the probability of a positive peak to occur within a repeat and then compare with the probability of occurring at any random position along the genome. Although some repeats might occur dispersed in the genome, only those inside cytological knob blocks (K180 and TR-1) or inside the centromere (CentC) were used for intersections.

In addition to the repeat families, we also performed the same analysis for the flanking regions of Transcription Start Sites (TSS), including an interval of 100 bp (50 bp downstream and upstream). The regions surrounding TSS are associated with hypersensitive MNase sites (Rodgers-Melnick et al. 2016). Therefore, while CentC was used as a reference of constitutive heterochromatin, the TSS regions were the reference of a highly accessible chromatin.

## RESULTS

### The knobs of maize inbred B73

Given the long-standing interest in maize heterochromatin and the tandem repeats known as knobs (McClintock 1930; Peacock et al. 1981), we set out to explore aspects of these cytological structures using newly available genome-wide chromatin sensitivity profiles available for a core set of different tissues for B73 during plant development (Turpin et al. 2018). For this study, we focused on knob repeat families as summarized in **Table 1**. Although the knobs have a similar cytological appearance, they are primarily made up of two different tandemly repeated DNA sequences, designated K180 for the abundant 180 bp satellite (Peacock et al. 1981) and the later described TR-1 for the 350 bp tandem repeat (Ananiev et al. 1998a). The maize knobs can be made up of primarily or exclusively one or the other or a mixture of both. For any given inbred line of maize, however, the position and cytological size of the knobs are stable and characteristic of that inbred (Longley 1939). This is typically referred to as “the knob composition”, which has been cataloged for a large number of maize inbred lines (Chughtai and Steffensen 1987; Chughtai and Steffensen 1989; Kato et al. 2004; Mondin et al. 2014).

**Table 1.**
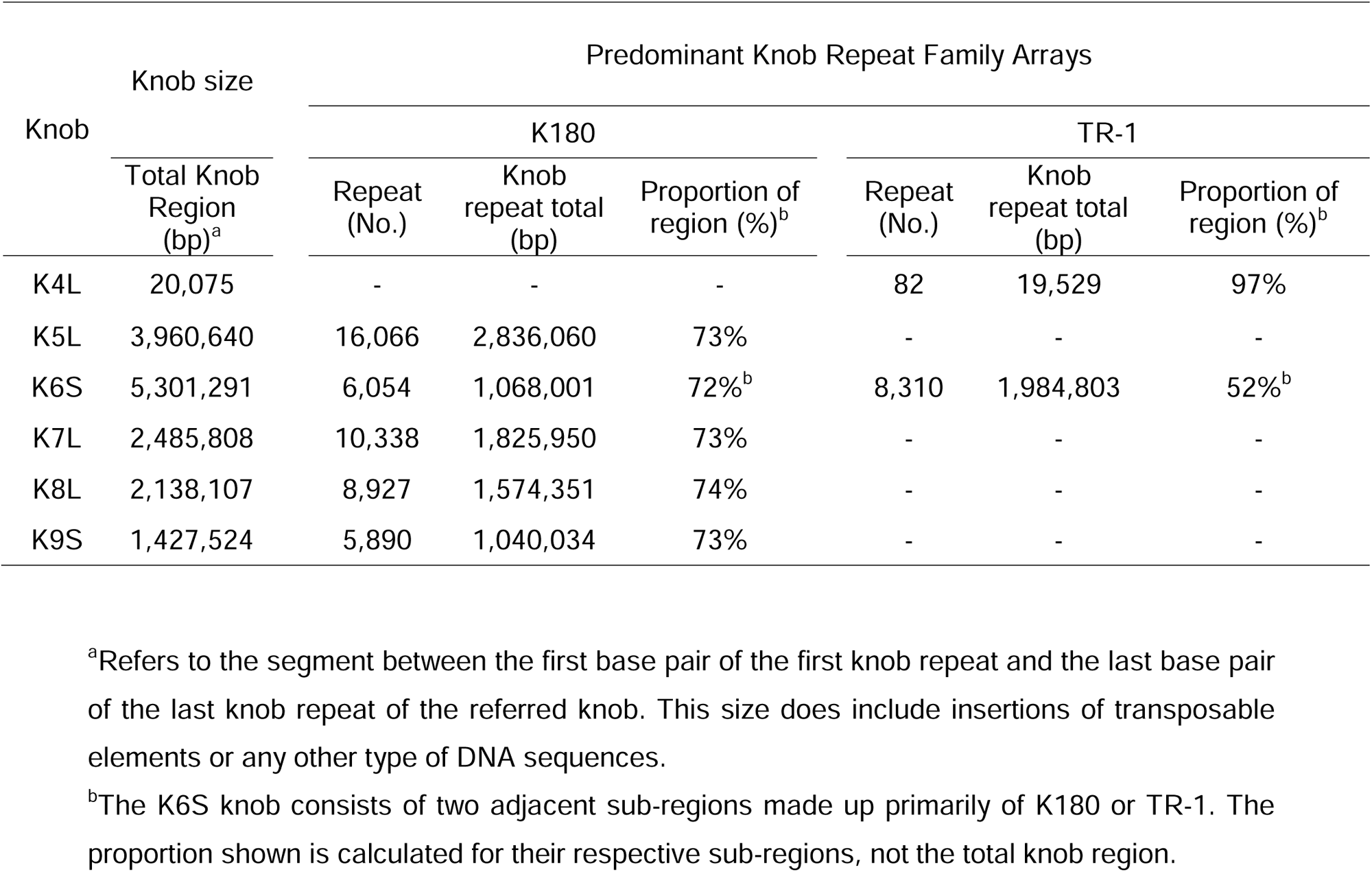
Size (bp) and composition of knobs in *Z. mays* B73v5 assembly.

The inbred used here, B73, is known from multiple studies to have several knobs (Kato et al. 2004; Ghaffari et al. 2013; Hufford et al. 2021) making it ideal to study knob behavior overall or individually. We filtered knob sequences according to their genomic EDTA annotation (Hufford et al. 2021) into K180 and TR-1, resulting in seven large arrays at six knob position loci (Table 1). Four of the large knobs are composed mainly of the K180 repeat family (K5L, K7L, K8L and K9S). These knobs, which range from ∼1.4 Mb to ∼4.0 Mb, have a very similar proportion of K180, from 72-74%. Other sequences interspersed with the knob region include TR-1, other repeat sequence families, and even a few genes. Another large knob, K6S, is made up of two adjacent sub-arrays, TR-1 and K180, with the TR-1 array being the distal and largest of the two (Table 1). One knob array, K4L, is quite short, only 20 kbp, below cytological visibility (Buckler et al., 1999), and composed mainly of TR-1 repeats. For the chromatin analysis, each of the seven knob arrays as defined by their predominant repeat type were analyzed as individual units.

### K180 but not TR-1 Knobs Exhibit Tissue-specific Variation in Chromatin Sensitivity Profiles

The multiple knobs together with chromatin profiling data provide an ideal situation to explore the chromatin state of knobs across the genome. For this, we visualized chromatin sensitivity from the differential nuclease sensitivity (DNS) profiles from Turpin et al. (2018), focusing on all seven knob arrays at all six knob loci as summarized in Figure 1. The DNS value (Vera et al. 2014) is derived from mathematical subtraction of the relative abundance of aligned library sequence coverages for light versus heavy MNase digests. The resulting DNS-seq coverage values are plotted along the genome as positive (blue) or negative (red) histogram tracks as shown in Figure 1. The positive/blue regions reflect the most easily cut and open/accessible chromatin, such as the promoter regions near the transcription start sites of genes. Conversely, the negative/red regions reflect more closed chromatin regions such as the protein-coding regions of genes, and the more inaccessible regions of the genome. Importantly, the digest sequence read coverage values were both read-depth and quantile normalized to allow for comparative analysis across the different digests and tissues.

**Figure 1.**
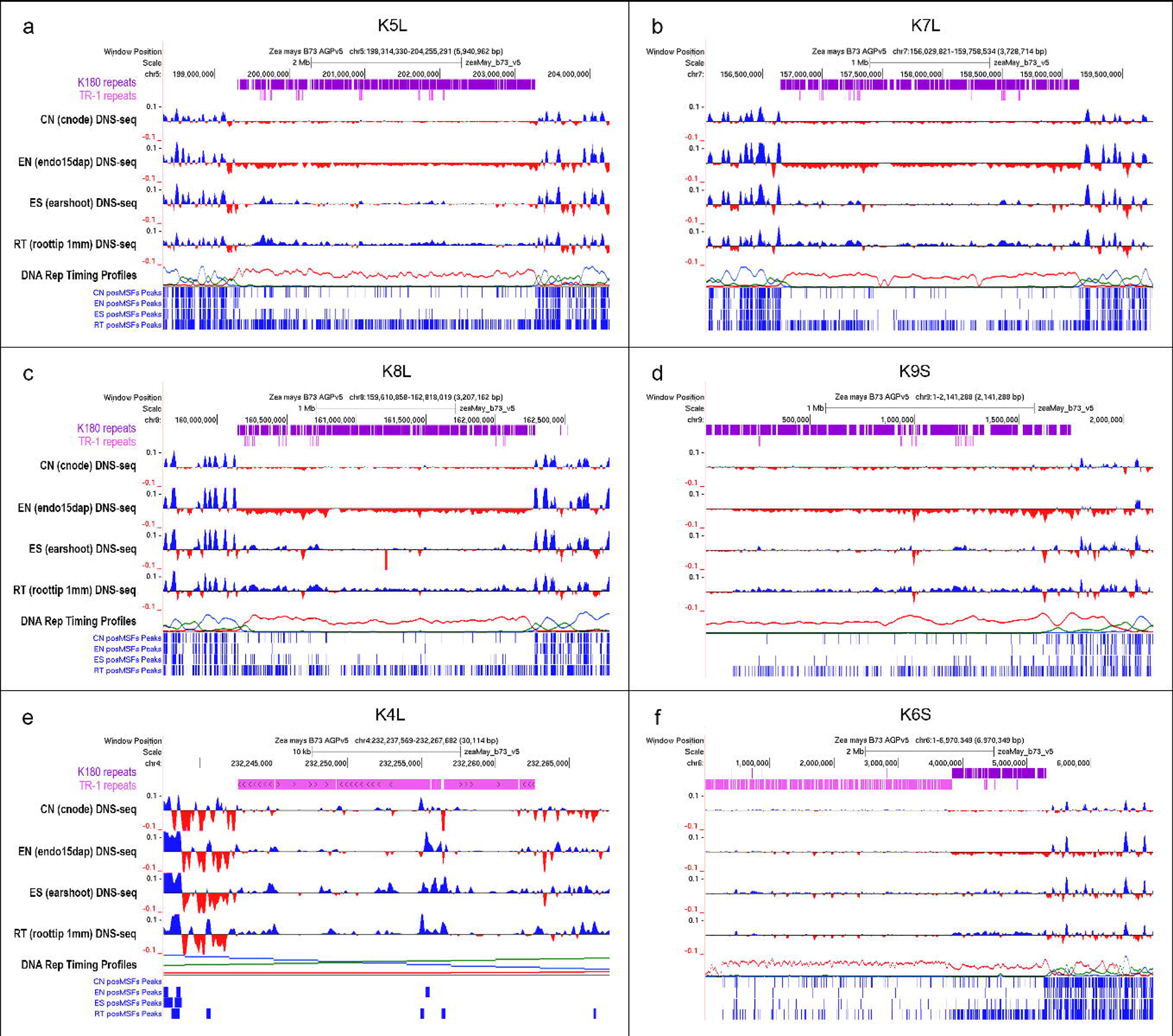
Chromatin accessibility patterns of knob regions in four different tissues of maize inbred B73. The regions corresponding to the K5L, K7L, K8L, K9S, K4L and K6S knobs are shown in (a), (b), (c), (d), (e) and (f) respectively. The tracks represent the knob repeat families: K180 (purple) and TR-1 (pink), which are above the DNS-seq profiles of four tissues (CN, EN, ES, and RT) and the DNA replication timing profiles track. The histogram tracks below correspond to the calling of positive peaks (in blue) using the MACS3 software (q=0.05). Scales in (a) and (f): 2Mb. Scales in (b), (c) and (d): 1Mb. Scale in (e): 10 Mb. DNS-seq profiles are scaled from −0.1 to 0.1.

The initial visual inspection of DNS-seq profiles on the genome browser revealed that TR-1 knobs had relatively more closed chromatin compared to the flanking non-knob chromatin, regardless of the tissue, as might be expected for heterochromatin (compare tissues for knobs in Fig 1a, b, c, and d). The K180 knobs were also generally more inaccessible than flanking non-knob chromatin, as expected. However, the K180 knobs were surprisingly varied in their sensitivity profile values when compared across the four different tissues. Specifically, the endosperm displayed the most inaccessible K180 knob chromatin (red regions in DNS-seq profile tracks, Fig 1a-d, f). The next most inaccessible K180 knob chromatin was seen for coleoptile nodes followed by the earshoots. Notably, the K180 knobs in root tip chromatin showed a contrasting pattern of more accessible K180 knob chromatin (blue regions in DNS-seq profile tracks, Fig 1a-d, f). Using the DNS-seq profile data, we called peaks with MACS3 for positive DNS-seq regions (bottom of panels in Fig. 1). The DNS-seq profiles and peak density both reflect the tendency for root tip K180 knob regions to present as more accessible compared to the three other tissue samples. This relative ranking of chromatin state across the four tissues seemed consistent for all five of the K180 arrays (Fig 1a-d, f). These observations, while striking, were primarily based on visual inspection of data patterns in the context of the genome browser viewing, prompting us to take a more statistical approach.

### K180 Knob Chromatin in Root Tips Exhibits a Significantly Open and a More Gene-like Chromatin State

To get a more quantitative assessment of the chromatin patterns, we employed a z-score-based approach to delve into the statistical significance of this tissue-specific variation as shown in **Table 2** and **Figure 2**. This allows for a comparison of observed versus expected chromatin states in regions of interest, such as K180 knobs, TR1-knobs, and other sequence types for comparison. For this, we investigated the DNS-positive open chromatin peak density in knobs relative to their observed density across the whole genome. In this analysis, a positive z-score indicates that the observed intersection of open chromatin peaks with knobs exceeds the expected value based on the mean of the randomized distribution, whereas a negative z-score indicates that it falls below the expected value. The magnitude of the z-score reflects how many standard deviations the observed value deviates from the mean of the randomized distribution.

**Figure 2.**
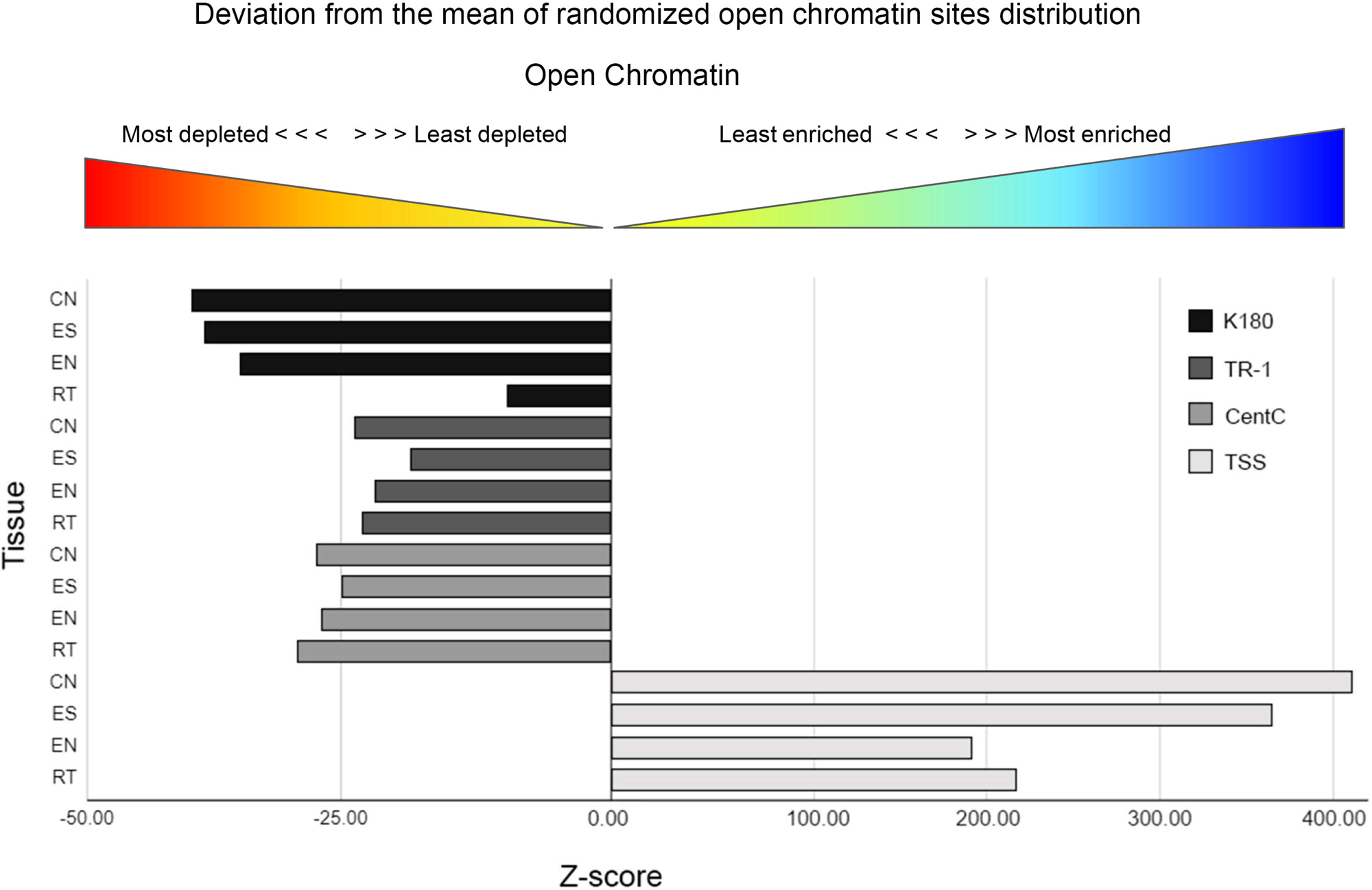
Graphical representation of the Z-score-based analysis for the enrichment/depletion of positive DNS-seq called peaks for different DNA sequences in four tissues of maize. More positive Z-score values indicate that the mean number of open chromatin (positive DNS peaks for MNase hypersensitive sites) co-locating with the target sequence is higher than expected based on the mean of the randomized distribution along the entire genome. The more negative Z-score values indicate a statistical depletion of open chromatin (positive DNS peaks) for each class of sequences. CN: Coleoptile Node; EN: Endosperm; ES: Earshoot; RT: Root Tip; TSS: Transcription Start Sites.

**Table 2.**
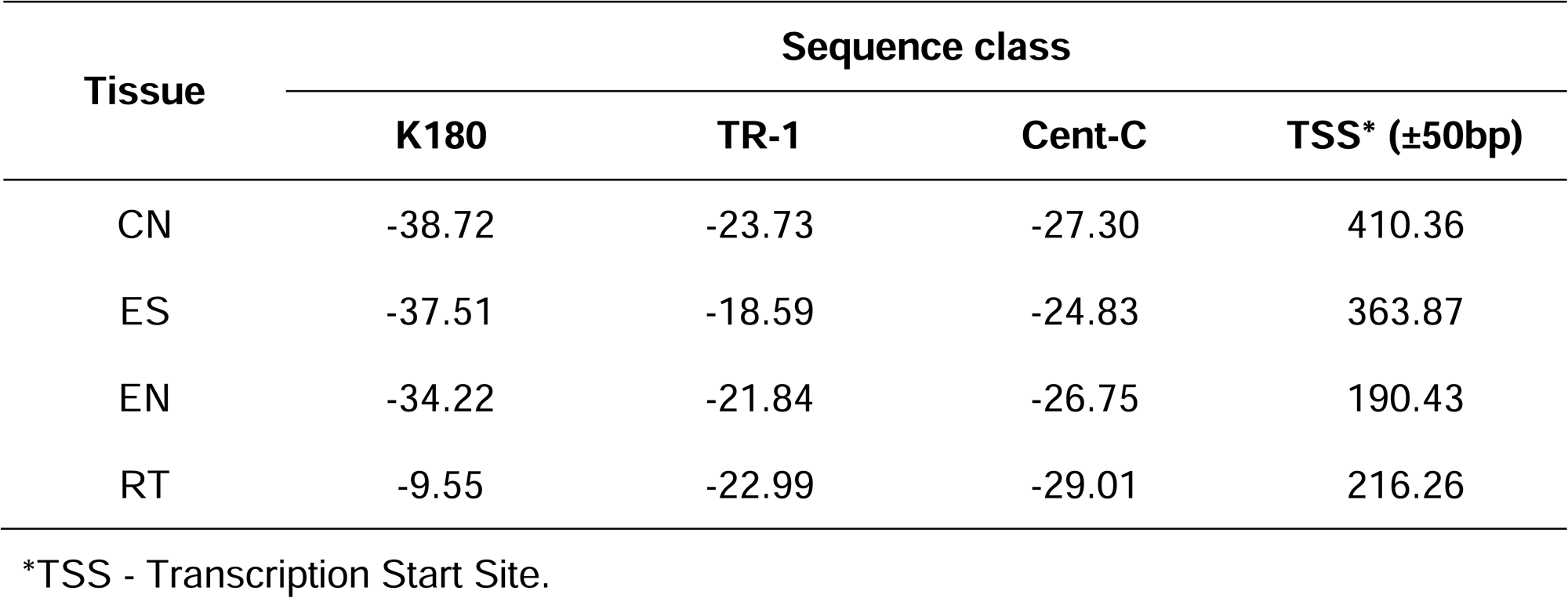
Z-score values for K180, TR-1, CentC and TSS in four tissues of *Zea mays* B73v5 obtained after intersection with positive called peaks from DNS-seq profiling data.

| Tissue | Sequence class |  |  |  |
| --- | --- | --- | --- | --- |
| | K180 | TR-1 | Cent-C | TSS* ($\pm 50$bp) |
| CN | -38.72 | -23.73 | -27.30 | 410.36 |
| ES | -37.51 | -18.59 | -24.83 | 363.87 |
| EN | -34.22 | -21.84 | -26.75 | 190.43 |
| RT | -9.55 | -22.99 | -29.01 | 216.26 |

\*TSS - Transcription Start Site.

The z-scores were calculated for four classes of sequences, three heterochromatic satellite DNA families, and a contrasting class of known open chromatin regions, 100 bp flanking the transcription start site (TSS) of the gene promoters. Two of these satellite DNA families are the main components of the knobs, comprising K180 and TR-1, which can occur separately or in combinations within individual knobs (Table 1). For instance, K5L, K7L, K8L, and K9S are made primarily of K180, K4L is made primarily of TR-1, and K6S is a composite of adjacent but non-overlapping blocks with the TR-1 array distal and the K180 array proximal to the centromere. The other two sequence classes were the CentC satellite DNA family, which forms constitutive heterochromatin not associated with knobs, and the TSS promoter regions of annotated genes, which represent open and highly accessible chromatin regions for comparison.

The results obtained from this chromatin state z-score-based approach concurred with the qualitative analysis. Specifically, the TR-1 knobs depicted fairly consistent chromatin state negative z-scores across the four tissues, ranging from −24 to −19. In striking contrast, the K180 knobs exhibited more tissue-specific variation. The root tip K180 knobs had a much less negative z-score, by several fold, for two types of comparisons. First, the K180 knob root tip (z-score = −10) is less negative than the K180 knobs in other tissues (z-scores of −38, −37, and −34). Similarly, even within the same root tip tissue, the K180 knobs (z-score = −10) were substantially more open than other heterochromatin regions including the TR-1 knob (z-score = −23) and the centromere-associated non-knob heterochromatin CentC (z-score of −29). This statistical analysis confirmed the significance of this tissue-specific variation in the chromatin state in knob repeats of K180, but not TR-1.

### Individuality of the Knobs in relation to chromatin accessibility

The previous analyses provided an overview considering all knobs composed of K180, TR-1, and their combinations, as well as centromeric regions and Transcriptional Start Sites. In this broader contextual analysis, the specific characteristics of each genomic region could not be observed. For instance, heterochromatic knobs exhibit significant variability in size and genomic complexity (Table 1). Therefore, to enable the visualization of potential individual features, each of these regions was examined in detail, and z-score values were analyzed and classified according to a heatmap. The objective of this analysis was to determine whether chromatin accessibility alterations in different plant tissues were restricted to knobs composed exclusively of K180, and whether the broader analysis presented earlier might have masked potential accessibility fluctuations in other heterochromatic regions.

Centromeric heterochromatin regions, represented by the Cent-C motif, exhibit high stability in chromatin accessibility patterns across nearly all chromosomes. The centromeres of chromosomes 2, 4, 5, 6, 8, 9, and 10 share the same chromatin accessibility pattern across all analyzed tissues (Fig. 3). Chromosome 3 displayed a slightly different pattern, but with low amplitude in z-score values. Moreover, no significant changes in z-score values were observed across different tissues in this chromosome. In contrast, chromosomes 1 and 7 exhibited distinct chromatin accessibility patterns in their centromeric regions compared to other chromosomes. The amplitude of z-score variation was 3.54 in chromosome 1 and 4.73 in chromosome 7. In both cases, the lowest z-score value was observed in the Ear Shoot, while the highest value was recorded in the Root Tip. Additionally, the z-score values were highly similar between the Endosperm and the Ear Shoot, as well as between the Coleoptile Node and the Root Tip. However, no statistically significant differences were detected between tissues.

**Figure 3.**
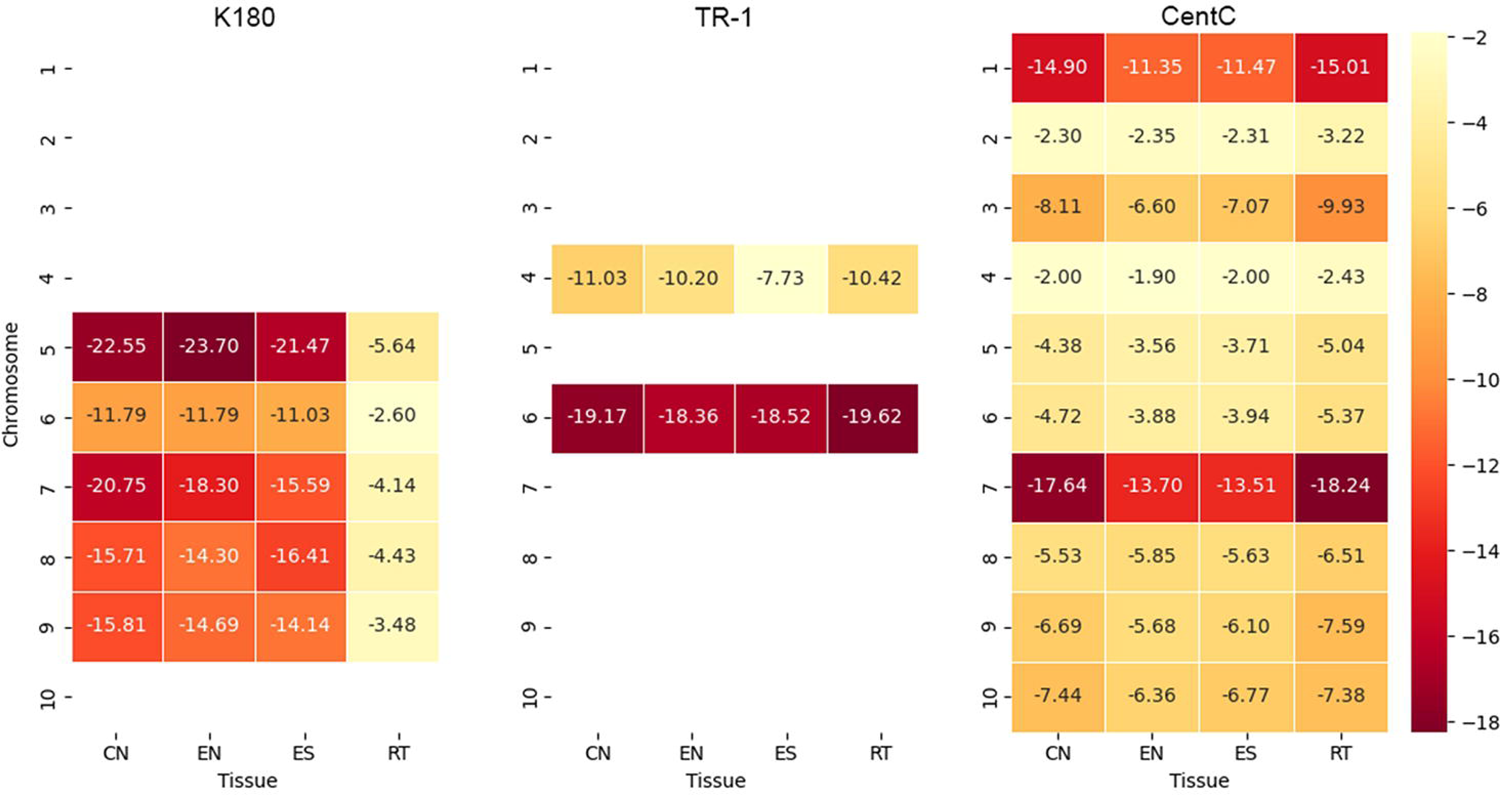
Chromosome by tissue comparison of individual Knob and CentC array chromatin states.

A similar trend of chromatin accessibility stability was observed for TR-1 knobs (Fig. 3). Both the TR-1 knob on the long arm of chromosome 4 and the one on the short arm of chromosome 6 exhibited no drastic variation in chromatin accessibility across different tissues. However, a notable difference was observed between these two regions from distinct chromosomes. The TR-1 knob on chromosome 4 presented lower z-score values and is a relatively small region, spanning approximately 20,000 base pairs and composed almost exclusively of the TR-1 motif (Table 1). Conversely, the TR-1 knob on chromosome 6, which exhibited high z-score values, corresponds to a much larger region containing nearly 1,000 times more repeats than the TR-1 knob on chromosome 4. Moreover, the TR-1 knob on chromosome 6 is juxtaposed to an extensive K180 region. These findings suggest that chromatin accessibility in this region may be associated with its size.

For K180 knobs, a significant shift in chromatin accessibility was observed between the Root Tip and other tissues. The largest amplitude of variation was detected in chromosome 5, with a difference of 18.06, while the smallest variation was found in chromosome 6, with a difference of 9.19 between the tissue with the highest value and the Root Tip, which consistently exhibited the lowest value. Additionally, no direct relationship was observed between tissue type and chromatin accessibility, except for the Root Tip. For example, the knob on chromosome 5 exhibited lower chromatin accessibility in the Endosperm, whereas the knob on chromosome 7 displayed reduced accessibility in the Coleoptile Node.

In this dataset, a clear distinction in z-score values was observed among the different knobs, excluding the Root Tip values. We found that these differences were directly correlated with the amount of K180 present in each knob and the total size of the knob region, except for chromosome 6. The knob on chromosome 5 exhibited the largest size, the highest number of K180 copies, and the highest z-score values, followed by chromosome 7, which ranked second in this regard. Although the knobs on chromosomes 8 and 9 did not differ significantly in z-score values, their distinction from the knobs on chromosomes 5 and 7 was evident. Regarding size and copy number, the knobs on chromosomes 8 and 9 were the smallest.

An exception was observed for the K180 knob on chromosome 6. Despite being structurally similar to the knob on chromosome 9, it is embedded in a distinct chromatin context. The total knob region in chromosome 6 comprises both K180 and TR-1, with K180 positioned more proximally and TR-1 more terminally. Interestingly, the TR-1 knob on this chromosome displayed z-score values comparable to those of the K180 knobs on chromosomes 5 and 7. Although the K180 knob on chromosome 6 shares structural similarities with those on chromosomes 7, 8, and 9, its z-score values were lower. We hypothesize that this discrepancy between the expected behavior of the K180 knob on chromosome 6 and the observed patterns in other knobs is due to its genomic and chromatin context. Notably, this knob region on chromosome 6 is located in close proximity to the nucleolus organizer region (NOR).

## DISCUSSION

The heterochromatic knobs of maize were first described by McClintock (1930, 1931) as occupying specific positions on chromosomes, with variation in their number and size generating unique cytological identities for traditional and commercial varieties, inbred lines, and hybrids (Longley, 1938; McClintock et al., 1981; Kato et al. 2004; Mondin et al., 2014; Carvalho et al., 2022). Initially, these heterochromatic structures were assumed to remain condensed throughout the entire cell cycle, with no apparent activity or dynamics, and were therefore considered gene-poor or genetically inert (Heinz, 1928). However, subsequent studies demonstrated that knobs undergo late replication during the cell cycle (Pryor et al., 1980; Bass et al., 2015), and alterations in this replication timing can lead to delays in their segregation due to incomplete separation of these heterochromatic blocks during anaphase, resulting in chromosomal breaks and the initiation of breakage–fusion–bridge (BFB) cycles (Fluminhan et al., 1996; Fluminhan et al., 1998; Santos-Serejo et al., 2018). From a genomic organization perspective, maize heterochromatic knobs are composed of two satellite DNA families. The first and most abundant is the K180 family (Peacock et al., 1981), followed by the TR-1 family (Ananiev et al., 1998a; Kato et al., 2004). These repetitive arrays are interspersed with transposable elements (Ananiev et al., 1998b). Although these two satellite DNA families can occur in an alternating pattern, their representation varies considerably among knobs. Some heterochromatic blocks are composed almost exclusively of K180 repeats—which are more frequent and abundant—while others exhibit a higher proportion of TR-1 repeats (Kato et al., 2004). Another striking feature is the strong correlation between knob number and abundance variability and the formation of clines with respect to latitude and altitude (McClintock et al., 1981; Porter and Rayburn, 1990). According to Buckler et al. (1999), these patterns indicate that knobs are not genetically neutral structures. Moreover, maize heterochromatic knobs are located in gene-rich regions, where they promote recombination suppression in their vicinity (Ghaffari et al., 2013).

Although maize heterochromatic knobs have been investigated from various perspectives, there are no reports in the literature that have examined the behavior of these structures throughout development. This gap may be attributed to the initial assumption that these regions remain inert throughout the entire cell cycle.

This study presents a detailed examination of the chromatin accessibility within maize knobs, revealing notable variation among four different tissues and two types of knob repeat families. Our findings show that the knobs composed of K180, but not TR-1, display tissue-specific variation in the chromatin accessibility profiles. Particularly, K180 knobs in root tips showed more open chromatin, contrasting sharply with the more closed states observed in the endosperm, ear shoot and coleoptile node. These results suggest an association between the repeat sequence composition and chromatin organization in relation to MNase sensitivity and provides new insights into the dynamic and complex nature of maize knob chromatin organization in different developmental contexts.

Naturally, it is not possible to make a direct comparison between chromatin accessibility in knobs and regions associated with gene activity, such as Transcriptional Start Sites (TSS), which exhibit very high levels of accessibility. Nevertheless, the observed differences in chromatin accessibility of knobs throughout development indicate that these regions display a previously unrecognized dynamic behavior. In contrast, the TR-1 knob on the long arm of chromosome 4 shows a remarkably stable chromatin accessibility pattern across all tissues analyzed. Meanwhile, the knob on the short arm of chromosome 6 contains two distinct domains—one composed predominantly of K180 and the other of TR-1. The K180 domain exhibits an oscillatory pattern similar to that observed in other knobs, with variability among root tips, coleoptile nodes, ear shoots, and endosperm. In contrast, the TR-1 domain displays no such variation, maintaining a stable pattern consistent with that of the TR-1 knob on chromosome 4L. This serves as an example of how chromatin accessibility behavior may be associated with repeat sequence type, as only the heterochromatic regions associated with K180 show oscillations in chromatin accessibility. Thus, it can be concluded that TR-1 exhibits a more stable pattern and a behavior more consistent with that expected of constitutive heterochromatin.

Typically, constitutive heterochromatin is associated with tightly compacted, transcriptionally inactive chromatin (Heitz, 1928). Therefore, the finding that K180 knobs in root tips exhibit a more open chromatin state compared to other tissues is intriguing. At first glance, one might relate this result to the high mitotic index of the root tips, which requires a more dynamic chromatin reorganization (Talbert et al., 2019). Nevertheless, the Z-score statistical approach revealed a significant deviation of K180 knob regions in root tips regarding the amount of accessible chromatin when compared with the rest of the genome. In addition, the same pattern did not occur to TR-1 knob repeat regions.

To confirm this observation, the constitutive heterochromatic region of the centromere—primarily associated with CentC—was also analyzed. These regions exhibited a consistent chromatin accessibility pattern across all tissues analyzed, characterized by a closed chromatin structure. Maize centromeric regions display a genomic organization similar to that of knobs, consisting of long arrays of CentC interspersed with transposable elements (Ananiev et al., 1998c; Nagaki et al., 2003; Birchler and Han, 2009; Piri et al., 2026). Nonetheless, the centromeres showed no changes in chromatin structural organization across the four tissues examined. These observations support our hypothesis of a direct association between the type of repetitive DNA and chromatin accessibility.

It is essential to discuss the robustness of the DNS-seq method used in this study. The differences observed in chromatin accessibility between K180 and TR-1 regions support the validity of the DNS-seq data and suggest that the variation in K180 accessibility is not a technical artifact. The uniform chromatin accessibility patterns observed for TR-1 and CentC repeats across all tissues corroborate with these findings. However, it is worth considering the technical challenges associated with sequencing repetitive regions, as repetitive DNA is known to pose difficulties in terms of accurate representation (Vera et al., 2014). Despite these challenges, the consistency of the results across biological replicates and tissues reinforces our confidence in the accuracy of the chromatin accessibility profiles reported here.

The tissue-specific variation in K180 knob chromatin accessibility challenges the traditional view of knobs as purely constitutive heterochromatin. This would imply a distinct epigenetic indexing of constitutive heterochromatin among different tissues. In the case of knobs, there appears to be an internal compartmentalization with respect to the distribution of histone modifications, as demonstrated by Shi and Dawe (2006). These authors showed that only H3K27me2 exhibited a specific and homogeneous signal within the knob, whereas H3K27me1 marked the periphery of the heterochromatic block, and H3K9me1 produced a weak signal. According to their findings, the H3K27me1 and H3K9me1 marks in maize cannot be strictly categorized as exclusive to either heterochromatin or euchromatin, as they are found in both. Given that all three marks are present to some extent in knobs, and that H3K27me1 is specifically localized to their periphery, it can be inferred that these cytological structures may contain more than one type of chromatin domain.

Considering that the knobs in the B73 maize inbred line exhibit significant differences in size and organization, we conducted an analysis that accounted for this individuality. Our results indicate that, although the overall developmental pattern of K180 knobs is generally consistent, individual knobs show variation in their chromatin accessibility dynamics throughout development. Notably, there is a lack of correlation between knob size and chromatin behavior. These findings are further supported by the individual analysis of centromeres and the separation of domains within the knob on the short arm of chromosome 6. Both TR-1 and CentC regions did not exhibit individualized patterns, instead maintaining a stable and uniform chromatin structure across all chromosomes analyzed. Therefore, once again, only knobs composed of K180 demonstrate such individualized chromatin behavior.

Previous research has indicated that heterochromatin can exhibit dynamic properties in response to environmental factors, such as stress (Dimitri et al., 2005; Pavet et al., 2006; Ting et al., 2011). The unique responsiveness of K180 has been previously reported by Hu et al. (2012). In that study, maize plants subjected to cold stress exhibited RNA expression derived from the K180 sequence. Although similar findings have not been reported in maize, the production of RNA from satellite DNA has been documented in other species. Notably, the work by Ting et al. (2011) demonstrated a substantial accumulation of such RNAs in various types of cancer. The exact reason for the production of RNA from satellite DNA located in constitutive heterochromatin regions under these conditions remains unclear. In our study, the focus was on the evaluation of tissues under typical environmental conditions. Therefore, the variation in K180 chromatin accessibility across tissues suggests a structural plasticity depending not only due to stress but also depending on the tissue context.

The tissue-specific accessibility of K180 repeats could have broader implications for genome organization and function. Knobs, previously regarded as cytological markers with limited functional roles, may be involved in regulating chromatin states during critical stages of development. For example, the more open chromatin in K180 knobs could allow for interactions with regulatory elements or chromatin remodeling factors, which could be essential for development. Moreover, the functional role of knobs in influencing phenotypic traits, such as flowering time (Carvalho et al., 2022), may be linked to their capacity to modulate chromatin states in a tissue-specific manner. From the perspective of individuality, these authors were able to correlate the presence or absence of the knob on the short arm of chromosome 9 with flowering time, while other knobs did not contribute to the trait. Taking these studies together raises the possibility that knobs contribute to the regulation of gene expression or chromatin architecture in ways that are more dynamic and context-dependent than previously recognized.

Future research should investigate the underlying mechanisms that govern the tissue-specific chromatin accessibility of K180 knobs. A comparative analysis of chromatin marks, such as histone modifications or DNA methylation patterns, in these regions could provide insights into the epigenetic regulation of knob chromatin. Additionally, transcriptomic studies could help determine whether the observed chromatin accessibility correlates with active transcription in root tips or other tissues. Given the emerging evidence of transcriptional activity within heterochromatic regions, including knobs, under stress conditions, it would be interesting to explore how chromatin accessibility in knobs responds to environmental cues. Furthermore, the role of knobs in genome stability and their potential interaction with chromatin remodeling complexes remain open areas for investigation.

Our study provides evidence for the tissue-specific modulation of chromatin accessibility in maize knobs heterochromatin, particularly within the K180 repeats. These findings suggest that knobs, long considered static and functionally inert, may have dynamic roles in genome regulation, with implications for development and genome organization. The results presented here reveal a new property of maize heterochromatic knobs, adding to previously described features such as late replication during the cell cycle, genomic organization, and epigenetic indexing. Given the extensive variability in composition and presence or absence of heterochromatic knobs across different maize genetic materials, our study provides a valuable reference for future investigations aiming to explore the behavior of these constitutive heterochromatin regions in greater detail.

## MAIN CONCLUSION

This study provides evidence for the tissue-specific modulation of chromatin accessibility in maize knobs heterochromatin, particularly within the K180 repeats.

## ACKNOWLEDGMENTS

MM is a PET-MEC fellowship. This work was supported by CNPq 408160/2021-7 and CNPq 404611/2024-9

## Notes

### Competing Interest Statement

The authors have declared no competing interest.

